# Active site mutations in the bacterial actin homolog FtsA impair direct interactions with the divisome in *Escherichia coli*

**DOI:** 10.1101/2022.02.25.482027

**Authors:** Josiah J. Morrison, Colby N. Ferreira, Evelyn M. Siler, Katie Nelson, Catherine E. Trebino, Benjamin Piraino, Jodi L. Camberg

## Abstract

During cell division in *Escherichia coli*, the highly conserved tubulin homolog FtsZ polymerizes and assembles into a ring-like structure, called the Z-ring, at the site of septation early in the division pathway. For recruitment to the membrane surface, FtsZ polymers directly interact with membrane-associated proteins. In *E. coli*, membrane recruitment and tethering of FtsZ are predominantly carried out by FtsA. FtsA shares structural homology with actin and, like actin, hydrolyzes ATP. Yeast actin detects nucleotide occupancy through a sensor region adjacent to the nucleotide binding site and adopts distinct conformations in monomeric and filamentous actin. Accordingly, bacterial actin homologs also display considerable conformational flexibility across different nucleotide-bound states and adopt a polymerized conformation. Here, we show that a cluster of amino acid residues in the central region of FtsA and proximal to the nucleotide binding site are critical for FtsA function in vitro and in vivo. Each of these residues are important for ATP hydrolysis, phospholipid (PL) binding, ATP-dependent vesicle remodeling, and recruitment to the divisome in vivo, to varying degrees. Notably, we observed that Ser 84 and Glu 14 are essential for ATP-dependent vesicle remodeling and magnesium-dependent membrane release of FtsA from vesicles in vitro, and these defects likely underlie the loss of function by FtsA(E14R) and FtsA(S84L) in vivo. Finally, we demonstrate that FtsA(A188V), which is associated with temperature-sensitive growth in vivo, is defective for rapid ATP hydrolysis and ATP-dependent remodeling of PL vesicles in vitro. Together, our results show that loss of nucleotide-dependent activities by FtsA, such as ATP hydrolysis, ATP-dependent PL vesicle remodeling, and membrane release, lead to failed Z-ring assembly and division defects in cells.

## Introduction

Bacterial cells divide by a widely conserved and essential process where multiple proteins collaborate to reshape and build new cell wall components, leading to the separation of a single parental cell into two separate progeny cells. This terminal stage of division is temporally and spatially coordinated within the cell to ensure that both progeny cells receive a copy of the bacterial chromosome and to ensure that the septum is established in the correct location at midcell. In *Escherichia coli*, the process of division and separation begins with the establishment of a network of cytoskeletal proteins, called the divisome, and the formation of a Z-ring. FtsZ polymers, containing protomers of the conserved, tubulin-like protein FtsZ, coalesce at midcell on the interior of the cell membrane, recruited there by an interaction with FtsA to form the Z-ring (Pichoff and Lutkenhaus 2005). FtsZ polymers are single stranded protofilaments that assemble head-to-tail by binding to GTP. FtsZ polymers dissociate in vitro as GTP is hydrolyzed, and dynamic assembly, disassembly and reassembly is thought to be a key activity that contributes to promoting the division event in vivo (McQuillen and Xiao 2020). FtsA, a bacterial cytoskeletal protein with homology to actin, recruits FtsZ to the phospholipid (PL) membrane and is also reported to dynamically assemble at the Z-ring along with FtsZ (Bisson-Filho et al., 2017, Caldas et al., 2019, Conti et al., 2018, Loose and Mitchison 2014).

FtsA is a widely conserved member of the Hsc70/sugar kinase/actin family and is present in most bacteria (Bork et al., 1992). *E. coli* FtsA assembles into actin-like polymers, promoted by ATP binding, (Morrison et al., 2022), and the C-terminal region of FtsA intercalates into the PL membrane bilayer (Conti et al., 2018, Pichoff and Lutkenhaus 2005, Yim et al., 2000). A truncated FtsA variant without the C-terminal 15 amino acid residues, FtsA(ΔMTS), polymerizes with ATP, indicating that PL binding is not required for polymerization; however, this variant weakly hydrolyzes ATP, in contrast to wild type FtsA’s highly processive ATPase activity in the presence of PL’s (Conti et al., 2018, Morrison et al., 2022). It is not understood how PL engagement coordinates or communicates with the active site of FtsA to regulate protein conformation or enzymatic activity, and further, how FtsA coordinates with FtsZ. When associated with PL vesicles, FtsA induces tubulation and vesicle remodeling when ATP is added, consistent with polymerization on the PL surface (Conti et al., 2018). When recruited to a supported lipid bilayer, FtsA assembly is surface restricted and can form PL-associated minirings that recruit FtsZ (Krupka et al., 2017). FtsA has also been shown to destabilize and remodel FtsZ polymers in vitro and one variant, FtsA(R286W), has been reported to require adenine nucleotide for FtsZ polymer destabilization (Beuria et al., 2009, Conti et al., 2018, Krupka et al., 2017).

To begin to address the interplay of ATP hydrolysis and function of FtsA, and protein communication between FtsA and FtsZ in the early steps of Z-ring assembly, we constructed mutations in the active site of FtsA and studied variant activity in vitro and in vivo. Specifically, we focused on residues near the predicted site of magnesium coordination since magnesium is expected to be required for catalysis and may therefore be important in regulating nucleotide-dependent conformational transitions. Using a variety of biochemical and biophysical assays, we compared ATP hydrolysis, PL-dependent remodeling, ATP-dependent lipid tubulation, and FtsZ polymer destabilization by FtsA wild type and mutant proteins and correlated these in vitro activities with function and ring localization in vivo. Our results show that amino acid residues near the catalytic site are important for supporting and regulating FtsA function.

## Material and Methods

### Bacterial strains and plasmids

Bacterial strains and plasmids used in this study are described in Table 1. Site-directed mutagenesis of *ftsA* was performed to construct substitution mutations S84L, A188V, D210A, and Y375A in pET-FtsA, pGfp-FtsA (pSEB293), and pQE9-FtsA (Morrison et al., 2022, Pichoff and Lutkenhaus 2005).

**Table 1.**
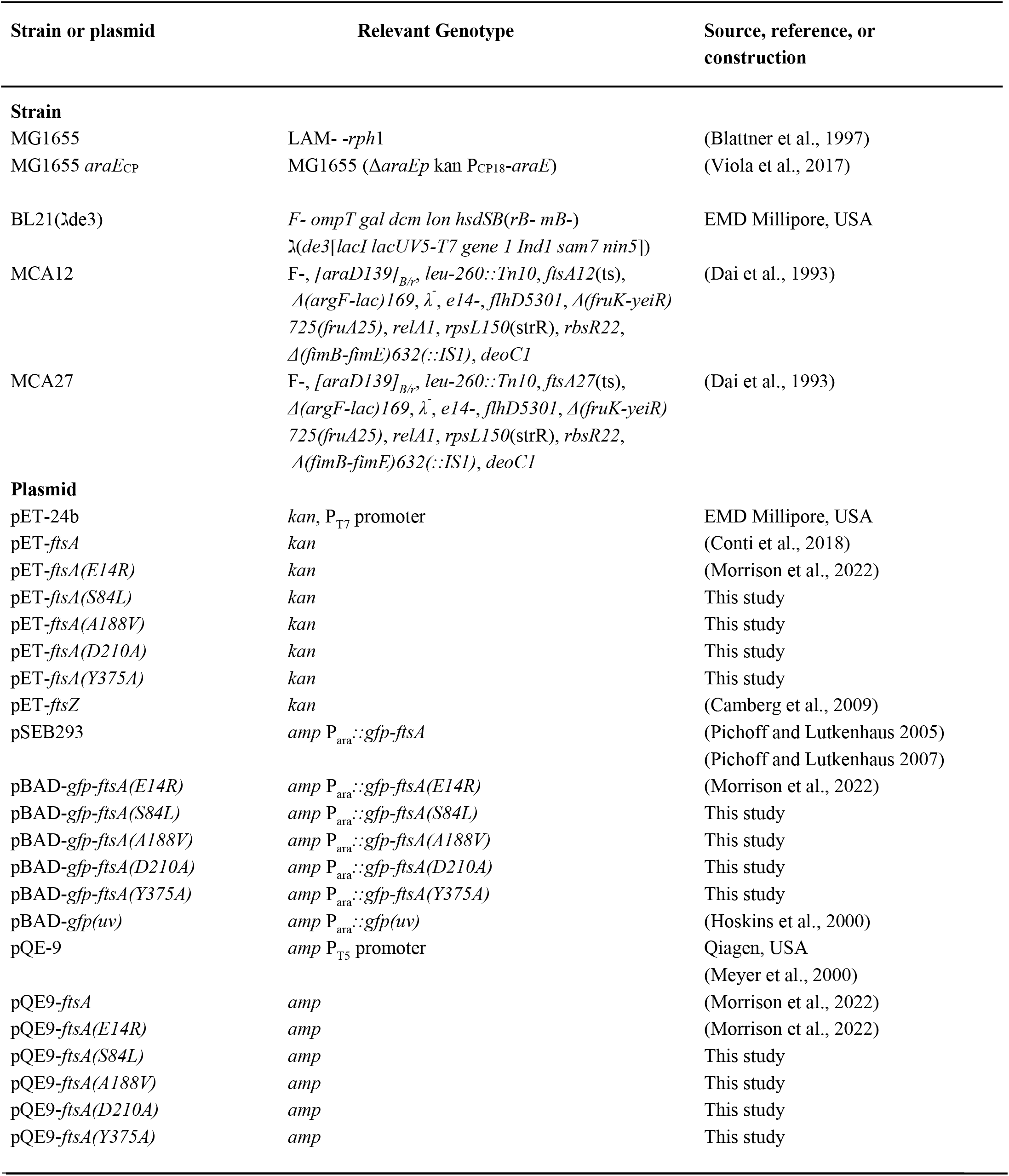
*E. coli* strains and plasmids.

### Structural Modeling

The amino acid sequence of *E. coli* FtsA was modeled onto the coordinates of *T. maritima* FtsA crystallized with magnesium and ATP (pdb: 1E4G) (van den Ent, 2000) by target-template alignment using ProMod3 by the Swiss-Model homology modeling server (Benkert et al., 2011, Bienert et al., 2017, Guex et al., 2009, Waterhouse et al., 2018). Distances were calculated in PyMol (v2.2.2).

### Temperature sensitive growth assays

*E. coli* strains containing expression plasmid pQE-9 (Qiagen), pQE9-*ftsA*, pQE9-*ftsA(E14R)*, pQE9*-ftsA(S84L)*, pQE9-*ftsA(A188V)*, pQE9-*ftsA(Y375A)*, or pQE9-*ftsA(D210A)* in temperature sensitive strains MCA27 [*ftsA*(S195P)] or MCA12 [*ftsA*(A188V)] were grown overnight in Lennox broth supplemented with 100 μg ml^-1^ ampicillin at 30 °C. The following day cultures were diluted into fresh LB with ampicillin to an OD_600_ of 0.05 and grown to an OD_600_ of 0.4 at 30 °C shaking (250 rpm). Cultures were diluted (2-log) into LB containing ampicillin and spotted (5 μl) onto LB agar plates with ampicillin and incubated at 30 °C (permissive) or 42 °C (restrictive) for 16 h.

### Protein purification

FtsA wild type and mutant proteins FtsA(E14R), FtsA(S84L), FtsA(A188V), FtsA(D210A), and FtsA(Y375A) were purified by ammonium sulfate fractionation, anion exchange, and size exclusion chromatography as described (Conti et al., 2018). FtsZ was purified as described (Camberg et al., 2009). Protein concentrations refer to FtsA monomers and FtsZ monomers.

### Phospholipid Assays

Phospholipid recruitment assays (25 μl) with FtsA wild type or mutant proteins containing substitutions (E14R), (A188V), (S84L) or (Y375A) were performed by incubating FtsA (wild type or mutant) proteins (1 μM) in 50 mM Tris pH 7.5, 150 mM KCl and 10 mM MgCl_2_ in the presence or absence of small unilamellar vesicles (SUV’s) (250 μg ml^-1^) with or without ATP (4 mM), or in the presence or absence of 25 mM EDTA where indicated, for 10 minutes at 30 °C followed by 21,000 x g centrifugation for 15 minutes at 23 °C. SUV’s were prepared from *E. coli* total membrane PL extracts (Avanti Polar Lipids) as described (Conti et al., 2018). Supernatants and pellets were analyzed by SDS-PAGE and coomassie staining. Percent of protein in the pellet fraction vs supernatant fraction was determined by densitometry using NIH ImageJ. Where indicated, lipid concentrations of FtsA fractions were determined by incubation of FtsA with FM 4-64FX (1.25 μg ml^-1^) (Invitrogen). Fluorescence intensity was measured with an Agilent Eclipse spectrofluorometer using an excitation wavelength of 565 nm and an emission wavelength of 745 nm (slit widths 5/10 nm). Concentration was determined by comparing to a PL standard curve.

Recruitment assays with FtsA and FtsZ were performed by incubating FtsZ (3 μM) with GTP (2 mM), in reaction buffer for 2 min, and then adding to a reaction containing FtsA wild type or mutant protein (2.5 μM), as indicated, pre-assembled with SUV’s (250 μg ml^-1^) and ATP (4 mM). Reactions were incubated for an additional 10 min at 30 °C, and then phospholipid vesicles were collected by low-speed centrifugation at 21,000 × *g* for 15 min. Supernatants and pellets were analyzed by SDS-PAGE and coomassie staining. Percent of protein in the supernatant fraction was determined by densitometry using NIH ImageJ.

To monitor ATP-dependent vesicle remodeling by light scatter, reaction mixtures (80 μl) containing reaction buffer and FtsA, FtsA(E14R), FtsA(S84L), FtsA(A188V), or FtsA(Y375A) (each at 1 μM) were monitored at 23 °C for 5 min to collect a baseline then ATP (4 mM) was added, where indicated, and reactions were monitored for an additional 15 min at on an Agilent Eclipse spectrofluorometer 450 nm with 5 nm slit widths.

### ATP Hydrolysis

ATP hydrolysis was measured by monitoring the amount of inorganic phosphate released at indicated temperatures in reactions (25 μl) using malachite green reagent (Enzo Life Sciences) for phosphate detection and comparing to a phosphate standard curve. ATP hydrolysis assays were performed at 37 °C unless otherwise indicated in reaction buffer containing 50 mM Tris (pH 7.5), 150 mM KCl, and 10 mM MgCl_2_, ATP (4 mM) and FtsA (1 μM) or FtsA mutants (1 μM). Where indicated, SUV’s (250 μg ml^-1^) were added to reactions.

### Transmission Electron Microscopy

Reaction buffer containing either FtsA, FtsA(E14R), FtsA(S84L), FtsA(A188V), or FtsA(Y375A) (each 4 μM), with or without ATP (4 mM) and SUV’s (250 μg ml^-1^), where indicated, were incubated for 10 min at 23 °C, applied to a 300-mesh carbon/formvar coated grid, fixed with glutaraldehyde (2.5%) and negatively stained with uranyl acetate (2%). Samples were imaged by transmission electron microscopy (TEM) using a JEM-2100 80 Kev instrument.

### FtsZ assembly assays

To collect FtsZ polymers by ultracentrifugation, reaction mixtures (25 μl) with FtsZ (3 μM) were prepared in reaction buffer containing 50 mM MES (pH 6.5), 100 mM KCl, and 10 mM MgCl_2_, 2 mM GTP and a nucleotide regenerating system containing acetyl phosphate (15 mM) and acetate kinase (25 μg ml^-1^). Where indicated, FtsA or FtsA mutant proteins (3 μM) and ATP (4 mM) were added. Reactions were incubated for 5 min at 23 °C and then centrifuged at 160,000 x *g* for 30 min in a Beckman TLA 120.1 rotor. Pellets and supernatants were resuspended in equal volumes, analyzed by SDS-PAGE and Coomassie staining, and quantified by densitometry using NIH ImageJ.

### Fluorescence Microscopy

Cultures of MG1655 *araE*_CP_ containing the plasmid pSEB293 (Pichoff and Lutkenhaus 2005) encoding GFP-FtsA, Gfp-FtsA(E14R), Gfp-FtsA(S84L), Gfp-FtsA(A188V), Gfp-FtsA(D210A), and Gfp-FtsA(Y375A) were grown overnight on solid LB Lennox media containing ampicillin (100 μg ml^-1^) at 30 °C and then streaked onto plates of the same composition and grown for 5 hours at 30 °C. Cells were collected and resuspended in 1x phosphate buffered saline (PBS) and, where indicated, incubated for 10 min in the dark at room temperature with FM 4-64FX (3 μg ml^-1^) (ThermoFisher). Cells were applied to a 5% agarose pad containing M9 minimal media with glucose (0.4%), and then a coverslip was added. Samples were visualized with a Zeiss LSM 700 confocal fluorescence microscope with excitation at 488nm and emission at 555nm for Gfp and 565nm and 744nm for FM 4-64FX respectively. Where indicated, a Nomarski prism was used to acquire differential interference contrast (DIC) images. All images were captured on an AxioCam digital camera with ZEN 2012 software. Cell lengths were measured using NIH ImageJ.

## Results

### Active site mutations in FtsA alter function in vivo, ATP hydrolysis, and PL binding

The FtsA active site is present at the center of the FtsA protomer, and accessibility to the central active site also appears to be maintained in actin-like filaments of *Thermotoga maritima* FtsA (Szwedziak et al., 2012, van den Ent and Lowe 2000). To investigate active site mutations in FtsA, we selected amino acid residues near the corresponding active site *magnesium* in the predicted structural model of *E. coli* FtsA (Fig. 1A) and performed site-directed mutagenesis to construct the following substitutions: E14R, S84L, A188V, D210A and Y375A. All selected amino acid side chains are approximately 3 to 5 angstroms (Å) from the magnesium cation modeled at the center of the *E. coli* FtsA active site (Fig. 1A and 1B) (Fig. S1A). Glu 14 (E14) and Asp 210 (D210) are situated in the phosphate-1 and phosphate-2 motifs, respectively (Bork et al., 1992), and thus predicted to be important for interacting with ATP phosphates and for hydrolysis activity. A recent report corroborated that FtsA(E14R) was defective for ATP hydrolysis in vitro and function in vivo (Morrison et al., 2022), providing a comparative basis to evaluate other mutations in this region of FtsA.

**Fig. 1.**
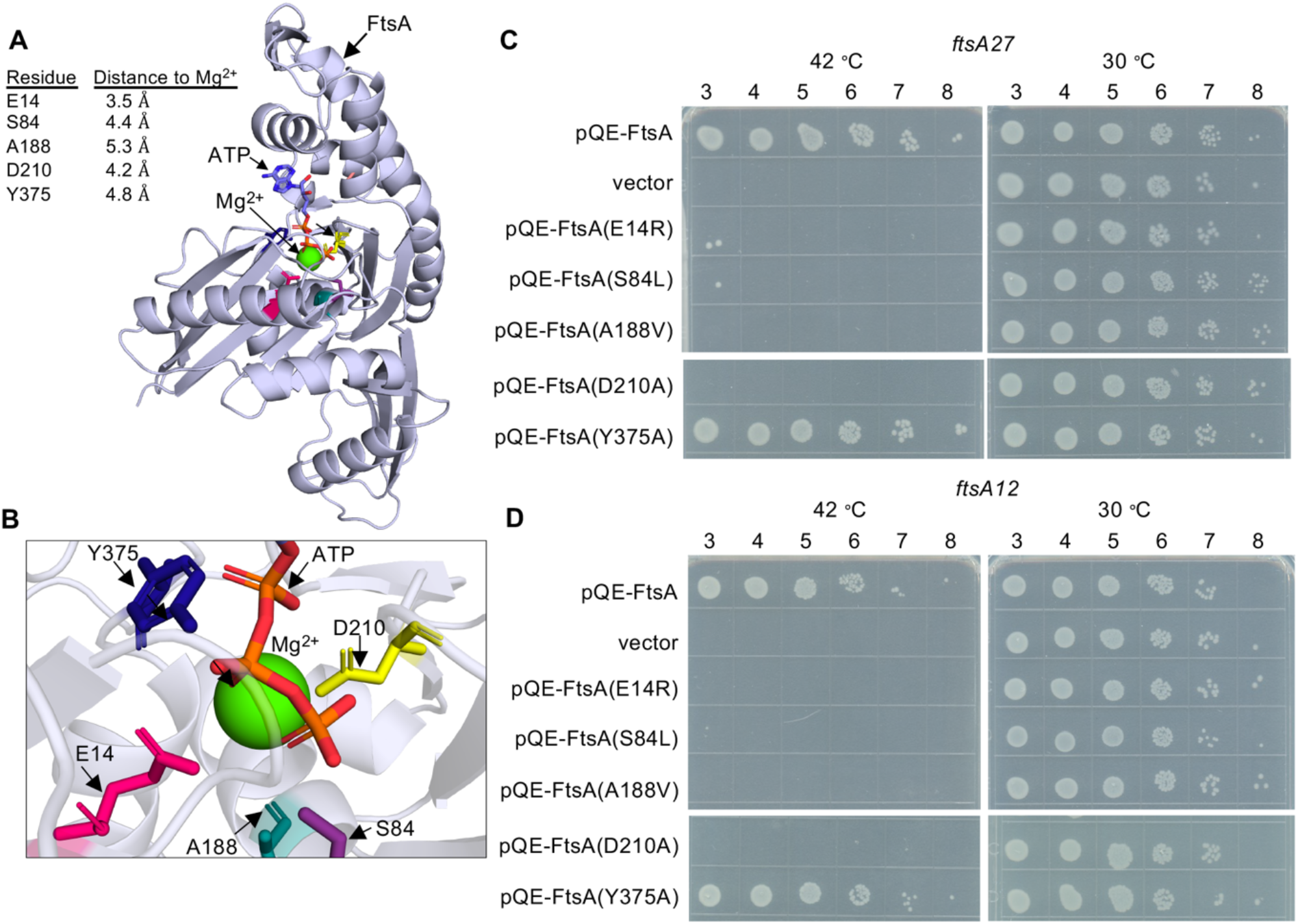
*E. coli* FtsA active site amino acid residues are important for function. (A) *E. coli* FtsA (residues 6-387) modeled onto *T. maritima* FtsA complexed with magnesium [green Corey-Pauling-Koltun (CPK) model] and ATP (stick) (pdb: 1E4G). *E. coli* amino acids Glu 14 (magenta), Ser 84 (purple), Ala 188 (teal), Asp 210 (yellow) and Tyr 375 (blue) are shown as sticks. Distances from modeled residues to Mg^2+^ are indicated in angstroms (Å). (B) Enhanced view of *E. coli* FtsA active site in (A). (C) Growth of *E. coli* strains containing expression plasmid pQE-9 (vector control), pQE9-*ftsA*, pQE9-*ftsA(E14R)*, pQE9*-ftsA(S84L)*, pQE9-*ftsA(A188V)*, pQE9-*ftsA(Y375A)*, or pQE9-*ftsA(D210A)* in MCA27 or (D) MCA12 temperature sensitive strains. Culture dilutions (2-log) were spotted (5 μl) on LB agar with ampicillin and grown as described in *Materials and Methods*. Plates were incubated for 16 h at 30 °C and 42 °C. Data shown is representative of three replicates.

Many *ftsA* substitution mutations located proximal to the FtsA ATP binding pocket have been reported to confer temperature-sensitive growth in *E. coli* (Fig. S1A), including the temperature-sensitive strains, MCA27 (*ftsA*27) and MCA12 (*ftsA*12) (Dai et al., 1993). MCA27 contains a substitution mutation in chromosomal *ftsA* converting Ser 195 to Pro, and MCA12 contains a substitution mutation that converts Ala 188 to Val. In both of these strains, growth is prevented at high temperature (42 ºC), but the cells grow robustly at the permissive temperature (30 ºC). To compare function of FtsA mutant proteins by a complementation assay, cultures of MCA12 and MCA27 cells containing a control vector (pQE-9) or an expression vector encoding wild type or mutant FtsA protein were cultured, spotted onto LB agar plates containing ampicillin, and then grown overnight at 30 and 42 ºC. As expected, cells containing pQE-FtsA grew at the restrictive temperature, whereas cells containing the control vector did not (Fig. 1C and 1D). Four of the FtsA mutant proteins were unable to complement in either background when expressed at the restrictive temperature, including FtsA(E14R), FtsA(S84L), FtsA(A188V), and FtsA(D210A) (Fig. 1C and 1D). FtsA(A188V) is a conservative mutation; however, it is the basis for the temperature-sensitive phenotype underlying MCA12. Our results demonstrate that increasing FtsA(A188V) expression from a plasmid is not sufficient to alleviate the temperature sensitive phenotype and suggest that the underlying defect may be related to a specific loss of FtsA function. Mutation of Ser 84 also led to loss of function, suggesting that this residue may provide an important contact or activity for FtsA. As expected, neither FtsA(E14R) nor FtsA(D210A) supported growth at the restrictive temperature in either strain background, consistent with previous reports of loss of function associated with mutation of these amino acids (Morrison et al., 2022, Pichoff and Lutkenhaus 2007, Pichoff et al., 2012, Yim et al., 2000).

To gain further insight into how magnesium site defects correlate with the various activities of FtsA that have been reported in vitro, we overexpressed and purified each FtsA mutant protein from *E. coli* and characterized it for the ability to hydrolyze ATP and bind to PL’s. FtsA(E14R), FtsA(S84L), FtsA(A188V), and FtsA(Y375A) were purified as described for wild type FtsA (Fig. S1B) (Conti, et al., 2018); however, we were unable to purify FtsA(D210A) and obtained no protein yield. Like wild type FtsA, all FtsA variants copurified with a small amount of PL’s, as has previously been described, although FtsA(A188V) copurified with significantly fewer PL’s (Fig. S1C) (Conti et al., 2018, Morrison et al., 2022). All FtsA variants were assayed for their ability to hydrolyze ATP, and all were observed to be partially defective for ATP hydrolysis under the conditions tested, with FtsA(A188V) being the most severe and retaining only 19% of wild type activity (Fig. 2A). FtsA(Y375A) was the least affected and retained 80% of wild type activity (Fig. 2A). In agreement with a previous report, we observed that FtsA(E14R) retains 61% of ATPase activity compared to wild type FtsA (Morrison et al., 2022), and FtsA(S84L) retained 68% of wild type ATPase activity (Fig. 2A). As FtsA(A188V) confers temperature-sensitive growth in vivo, we compared FtsA(A188V) ATP hydrolysis in vitro at 30 °C and 42 °C, which corresponds to the restrictive temperature for growth in vivo (Dai et al., 1993). However, we observed no further defect, or stimulation, of the rate of ATP hydrolysis by FtsA(A188V) as temperature increased and under the conditions tested (Fig. S2). The results suggest that the underlying cause of temperature-sensitive growth is not directly related to FtsA ATP hydrolysis rates.

**Fig. 2.**
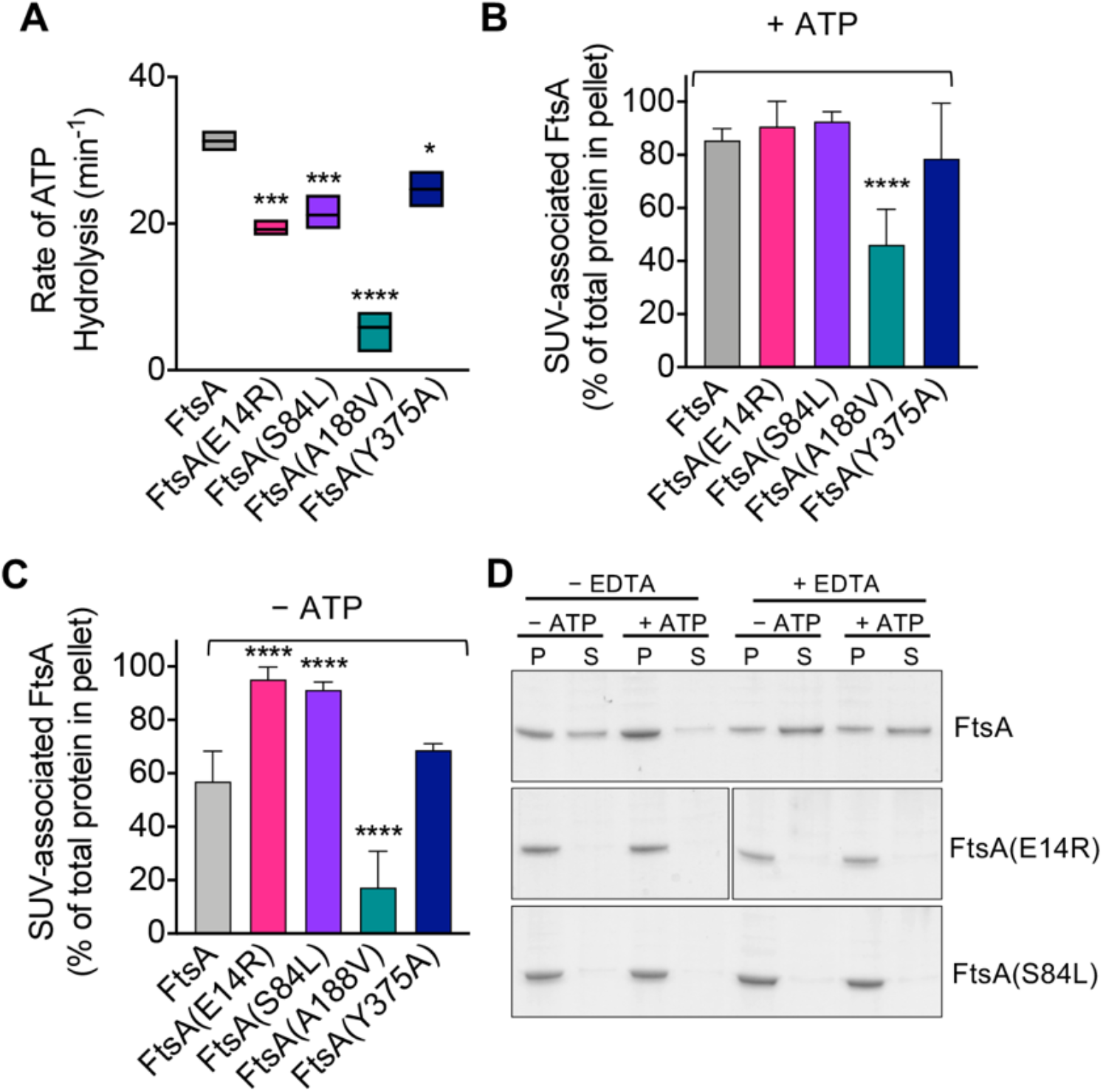
ATP hydrolysis and PL binding activity of FtsA wild type and mutant proteins. (A) ATP hydrolysis rates of FtsA wild type and FtsA mutant proteins (1 μM) were assayed by measuring the release of phosphate over time in reactions containing ATP (4 mM) as described in *Materials and Methods*. Statistical analyses for data in (A) performed by comparison to FtsA wild type value (*, p-value < 0.02; ***, p-value < 0.001; ****, p-value < 0.0001). (B) Recruitment of FtsA wild type and mutant proteins (1 μM) in assay buffer supplemented with SUV’s (250 μg ml^-1^) with and without (C) ATP (4 mM) as described in *Materials and Methods*. Data shown is an average of at least three replicates with error represented as S.E.M. Statistical analyses for data in (B) and (C) performed by comparison to FtsA wild type values (****, p-value < 0.0001). (D) Phospholipid recruitment of FtsA, FtsA(E14R), and FtsA(S84L) (2 μM) to SUV’s (250 μg ml^-1^) in the presence and absence of ATP (4 mM) and EDTA (25 mM) to chelate Mg^2+^ as described in *Materials and Methods*. Data shown is representative of three replicates.

Next, to determine if FtsA mutant proteins are defective for ATP hydrolysis due to a failure to bind PL’s, which is essential for rapid ATP turnover by wild type FtsA (Conti et al., 2018), we performed a PL recruitment assay by low-speed centrifugation in the presence of exogenous SUV’s (250 μg ml^-1^), which were added in excess to reactions to supplement copurifying PL’s (Fig. S1C), and then we collected PL associated proteins (Fig. 2B and 2C). We observed that all FtsA mutant proteins fractionated with the PL pellet in the presence of ATP, and that only FtsA(A188V) was partially impaired for PL recruitment (Fig. 2B). In the absence of ATP, 29% less wild type FtsA fractionated with the PL pellet; however, FtsA(E14R) and FtsA(S84L) remained robustly associated with the PL fraction (Fig. 2C). In contrast to FtsA(E14R) and FtsA(S84L), FtsA(A188V) was poorly recruited to PL’s when ATP was omitted (Fig. 2C), although it was recruited to a lesser extent than with ATP (Fig. 2B). These results show that FtsA(A188V) is defective for binding to PL’s, compared to wild type FtsA, and that FtsA(E14R) and FtsA(S84L) bind to PL’s promiscuously without ATP.

FtsA(S84L) and FtsA(E14R) are partially defective for ATP hydrolysis (Fig. 2A), showing 32% and 39% reduced rates of ATP hydrolysis compared to wild type FtsA, respectively. However, in the PL recruitment assay, both FtsA mutant proteins associated with PL’s similarly in the absence and presence of ATP, in contrast to wild type FtsA, which displays reduced PL association without ATP. One possibility for this is that FtsA(E14R) and FtsA(S84L) may persist in the PL-bound conformation regardless of nucleotide status. Glu 14 and Ser 84 are close to the predicted site of magnesium binding in FtsA (3.5 Å and 4.4 Å, respectively) (Fig. 1A and 1B), therefore we hypothesized that magnesium occupancy at this site is involved in the conformational transitions between PL-associated and PL-released FtsA. To investigate this further, we repeated the PL recruitment assay with FtsA and ATP in the presence of EDTA to chelate the magnesium and observed that without magnesium 60% of FtsA remained soluble and not bound to PL’s when ATP was present, whereas less than 5% remained soluble and not bound to PL’s when ATP and magnesium were available (Fig. 2D). This suggests that Mg^2+^ *and* ATP occupancy promote PL-binding by FtsA. Next, we tested if FtsA(E14R) and FtsA(S84L) were also released from PL’s when treated with EDTA but observed that both proteins remained associated with PL’s under all conditions tested (Fig. 2D). This suggests that mutations in Glu 14 and Ser 84 disrupt the ability to support release from the membrane, suggesting that in the apo state, without Mg^2+^ and ATP, the FtsA mutant proteins persist in a PL-bound conformation in vitro.

In previously reported light scattering assays (Conti et al., 2018, Morrison et al., 2022), FtsA was shown to remodel and tubulate PL vesicles when ATP is added to the reaction, and this activity likely corresponds to ATP-dependent polymerization in vitro (Morrison et al., 2022). To determine if FtsA mutant proteins are competent for tubulation of vesicles, we monitored 90° light scatter of reactions containing FtsA wild type and mutant proteins, and then added ATP to determine if ATP induces tubulation. In the absence of ATP, the light scatter signal produced by FtsA remained constant. We observed that wild type FtsA and FtsA(Y375A) both had a large increase in light scatter after ATP was added to the reactions (Fig. 3); however, FtsA(E14R), FtsA(S84L), and, to a lesser extent, FtsA(A188V) were impaired for vesicle reorganization. The results suggest that while FtsA and FtsA(Y375A) are competent for ATP-induced vesicle remodeling, FtsA(E14R), FtsA(S84L) and FtsA(A188V) are defective.

**Fig. 3.**
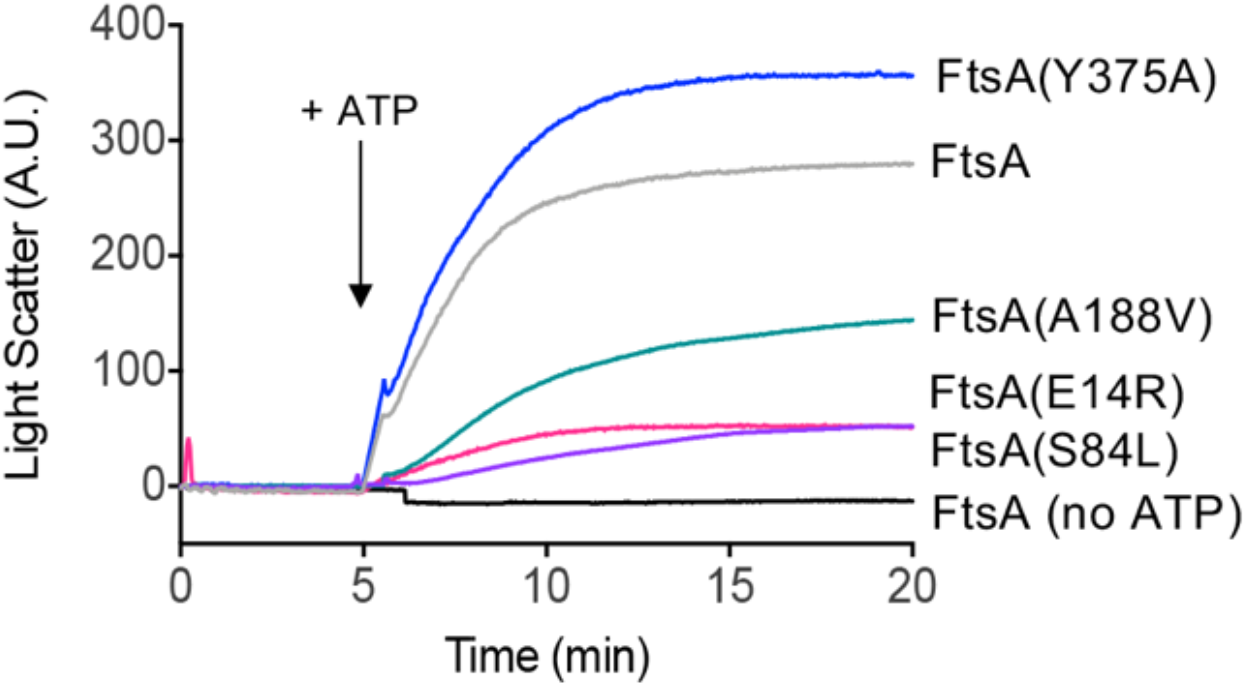
ATP-dependent PL remodeling by FtsA wild type and mutant proteins. FtsA wild type and FtsA mutant proteins (1 μM) were monitored by 90º angle light scatter at 450 nm. A baseline signal was measured for 5 min, ATP (4 mM) was added, and reactions were monitored for an additional 15 min. Data shown is representative of three replicates.

Finally, we visualized FtsA wild type and mutant proteins by negative stain transmission electron microscopy (TEM) with and without ATP and supplemental SUV’s to detect PL tubulation. We observed ATP-stimulated tubule formation with wild FtsA, FtsA(Y375A) and, to a lesser extent, FtsA(A188V), but failed to detect tubulation by FtsA(E14R) or FtsA(S84L) (Fig. 4). The tubules observed were approximately 50-80 nm in length. These results are consistent with the reduced light scatter observed after addition of ATP by FtsA(E14R), FtsA(S84L), and to a lesser extent, FtsA(A188V), compared to wild type FtsA (Fig. 3).

**Fig. 4.**
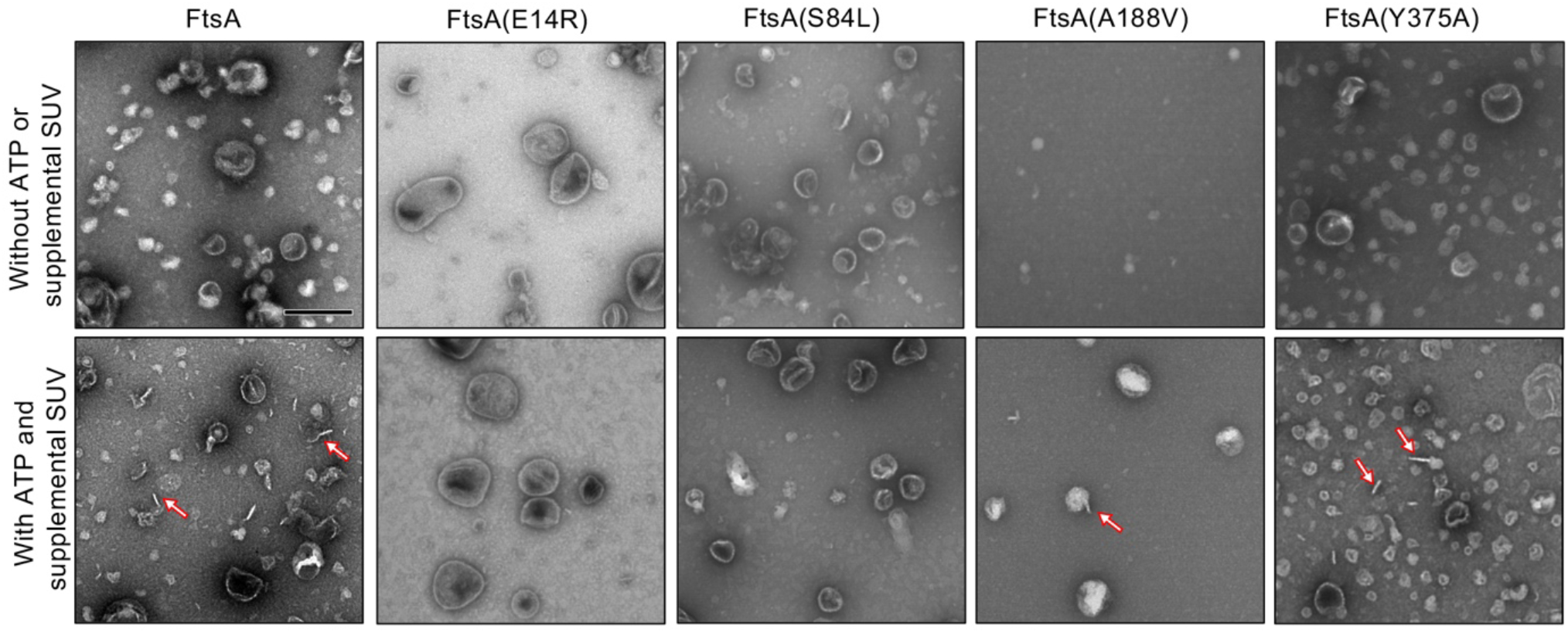
FtsA promotes vesicle tubulation with ATP. FtsA wild type and mutant proteins, FtsA(E14R), FtsA(S84L), FtsA(A188V), FtsA(Y375A), (4 μM) were incubated in the presence and absence of ATP (4 mM) and SUV’s (250 μg ml^-1^) and visualized by TEM. Arrows indicate tubulated liposomes. Scale bar is 200 nm.

### FtsA mutant proteins are defective for destabilizing FtsZ polymers in vitro

FtsA binds to FtsZ polymers, recruits the polymers to PL’s, and promotes destabilization of polymerized FtsZ, releasing it into the soluble fraction while FtsA remains associated with PL’s (Conti et al., 2018, Morrison et al., 2022). To determine if FtsA mutant proteins are defective for disassembly of FtsZ polymers, we assembled FtsZ polymers with GTP, with and without FtsA and ATP, and then quantified soluble and polymerized fractions of FtsZ by SDS-PAGE (Fig. 5A) (Fig. S3). We observed that without FtsA and ATP, 29.3 ± 2.4% of FtsZ, after incubation with GTP to induce polymerization, remained in the soluble, non-polymerized fraction after high-speed ultracentrifugation, and the majority of FtsZ sedimented as polymers (Fig. 5A) (Fig. S3). However, in the presence of FtsA and ATP, 43.3 ± 1.6% of FtsZ localized to the soluble fraction, suggesting that FtsA destabilizes FtsZ polymers and reduces the amount of FtsZ polymers collected by ultracentrifugation, consistent with destabilization or antagonizing activity by FtsA reported in other studies (Conti et al., 2018, Krupka et al., 2017, Morrison et al., 2022). The amount of FtsZ detected in the supernatant after incubation with FtsA(Y375A) was similar to wild type FtsA (39.8 ± 4.4%) (Fig. 5A). However, we detected no increase in the amount of FtsZ detected in the supernatant after incubation with FtsA(E14R), FtsA(S84L), or FtsA(A188V), suggesting that the FtsA mutant proteins are defective for destabilizing FtsZ polymers under the conditions tested (Fig. 5A).

**Fig. 5.**
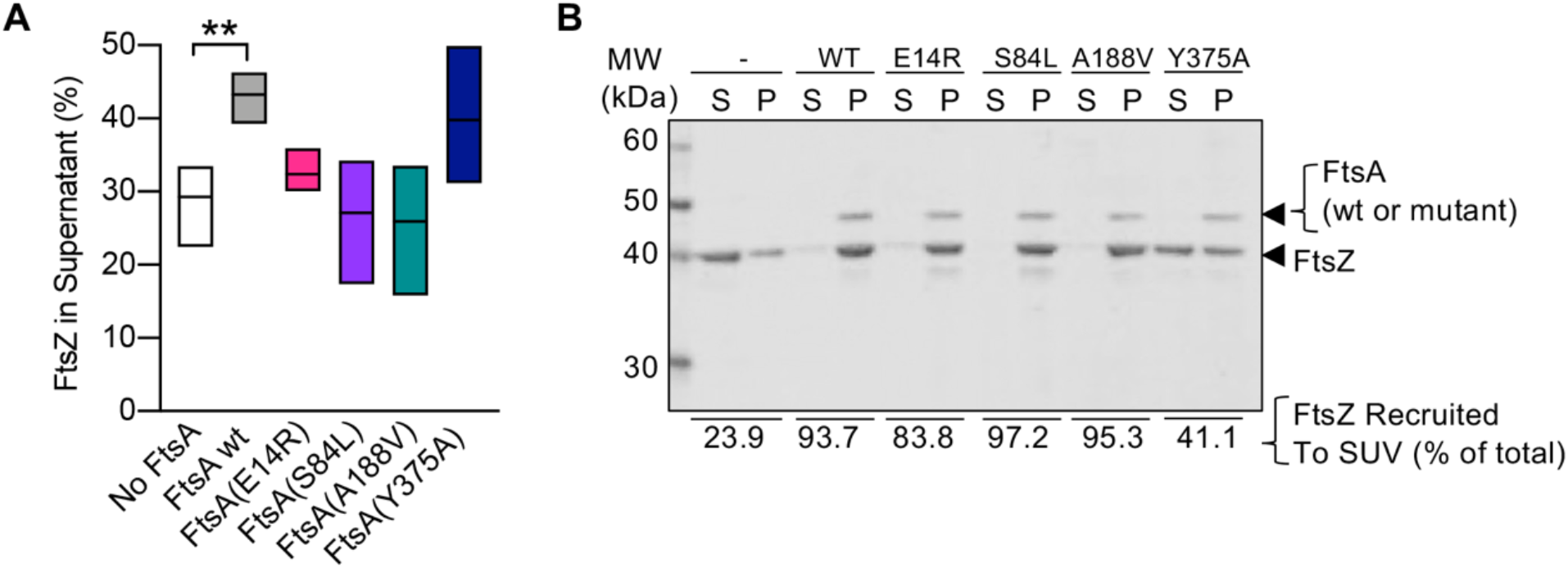
FtsA mutant proteins fail to destabilize FtsZ polymers. (A) Reactions containing FtsZ (3 μM) and, where indicated, GTP (2 mM), FtsA, FtsA(E14R), FtsA(S84L), FtsA(A188V), FtsA(Y375A) (3 μM), and with ATP (4 mM) and a regenerating system containing acetyl phosphate (15 mM) and acetate kinase (25 μg ml^-1^) were incubated for 5 min then fractionated by ultracentrifugation. Pellets and supernatants were visualized by SDS-PAGE and coomassie staining and quantified by densitometry. Data shown is an average of at least three replicates with error represented as S.E.M. Statistical analyses for data in (A) performed by comparison to ‘No FtsA’ (**, p-value = 0.0026). (B) Assays to recruit FtsZ to PL through an interaction with FtsA were performed by incubating FtsZ (3 μM) with GTP (2 mM) and then adding to a reaction containing FtsA wild type or mutant protein (2.5 μM), SUV’s (250 μg ml^-1^), and ATP (4 mM) as described in *Materials and Methods*. SUV’s and bound proteins were collected by centrifugation and analyzed by SDS-PAGE. Percent of FtsZ recruited to SUV’s is shown. Data shown is representative of three replicates.

Next, to determine if the failure to destabilize FtsZ polymers resulted from an inability of FtsA mutant proteins to interact with FtsZ, we performed a PL recruitment assay. In this assay, FtsZ polymers, assembled with GTP, were incubated with FtsA, ATP, and SUV’s, and then the SUV’s and bound proteins were collected by low-speed centrifugation and analyzed by SDS-PAGE. We observed that all FtsA mutant proteins were capable of recruiting FtsZ to varying degrees, including FtsA(S84L) and FtsA(A188V), and, both to a lesser extent, FtsA(E14R) and FtsA(Y375A) (Fig. 5B). Together, these results suggest that FtsA mutant proteins FtsA(S84L) and FtsA(A188V), like FtsA(E14R), bind to FtsZ but are defective for destabilizing FtsZ polymers in vitro.

### FtsA mutant proteins are defective for ring assembly in vivo

To determine if FtsA mutant proteins that are defective for function in vivo in temperature-sensitive growth assays (Fig. 1C and 1D) are also defective for localizing to a ring during division, we constructed and expressed each FtsA mutant protein as a fluorescent fusion protein to Gfp (green fluorescent protein) and imaged live dividing cells by fluorescence microscopy. We observed that expression of Gfp-FtsA from a plasmid [pSEB293; (Pichoff and Lutkenhaus 2005)] led to the visualization of clear, robust rings (A-rings) at midcell in 51.5% of the population, and cells had a mean cell length of 2.36 ± 0.05 μm (Fig. 6A and 6B). We also observed A-rings in cells expressing Gfp-FtsA(Y375A) (17.5%) and Gfp-FtsA(A188V) (15.0%), indicating that both mutant proteins are partially defective for A-ring formation. Cells expressing Gfp-FtsA(S84L), FtsA(D210A), or FtsA(E14R) had few to no visible rings (6.9%, 0% and 0%, respectively), and a mean cell length similar to cells expressing Gfp-FtsA (2.51 ± 0.07 μm, 2.23 ± 0.05 μm, and 2.48 ± 0.06 μm, respectively) (Fig. 6A and 6B). However, cells expressing Gfp-FtsA(A188V) and Gfp-FtsA(E14R) were shorter than cells expressing Gfp-FtsA, with mean cell lengths of 1.92 ± 0.08 μm and 1.87 ± 0.05 μm, respectively. These results show that FtsA mutant proteins that are most defective for restoring growth of temperature-sensitive strains at the restrictive temperature [i.e., Gfp-FtsA(D210A), Gfp-FtsA(E14R), Gfp-FtsA(S84L), and Gfp-FtsA(A188V)] (Fig. 1C and 1D) are also most defective for localizing to rings during division.

**Fig. 6.**
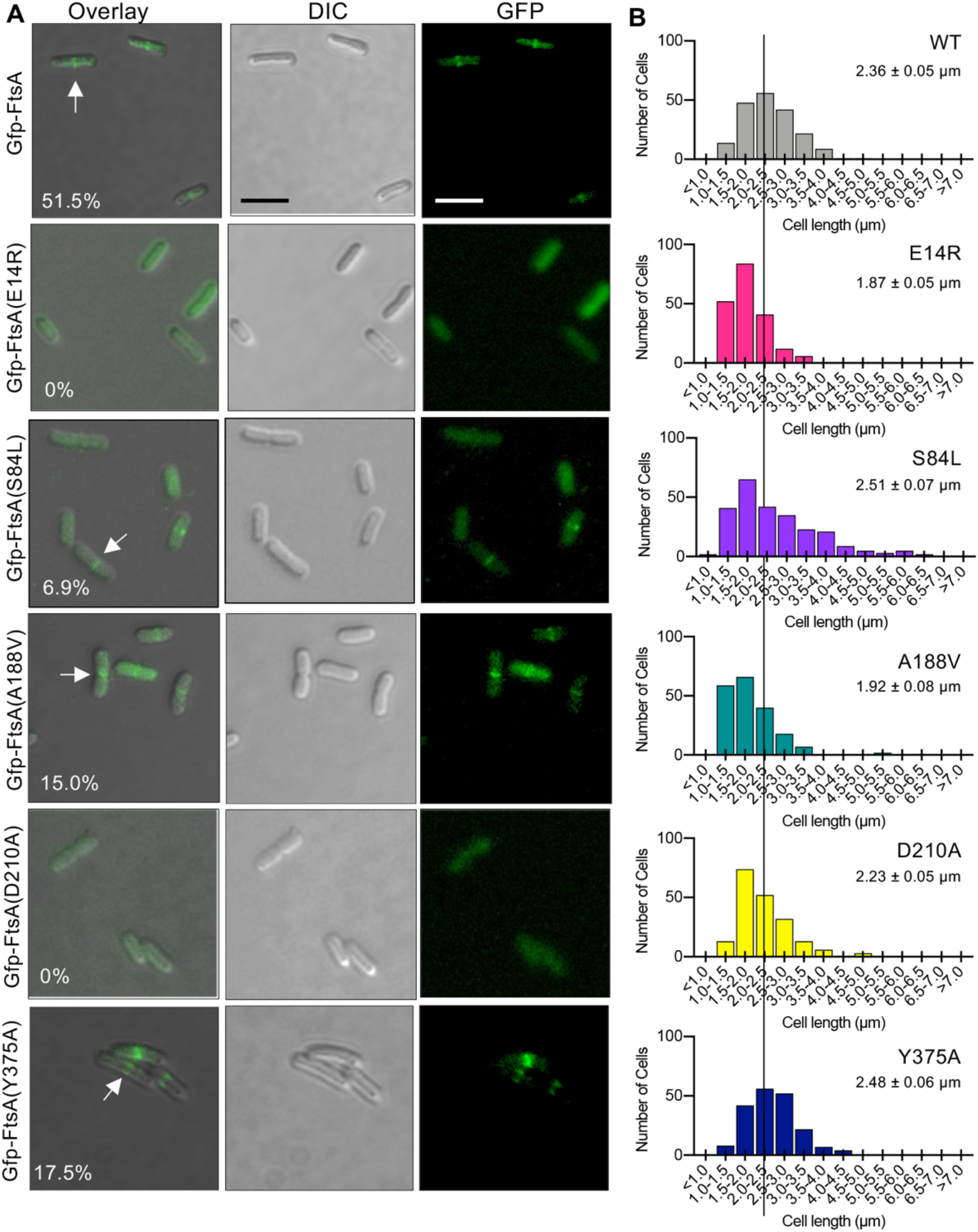
Fluorescence microscopy of Gfp-tagged FtsA mutant proteins. (A) Confocal fluorescence and differential interference contrast (DIC) microscopy of *E. coli* MG1655 *araE*_*CP*_ cells expressing plasmid-encoded Gfp-FtsA, Gfp-FtsA(E14R), Gfp-FtsA(S84L), Gfp-FtsA(A188V), Gfp-FtsA(D210A), or Gfp-FtsA(Y375A). Cells were grown as described in *Materials and Methods*. Scale bars are 2 μm. White arrows indicate A-rings. (B) Plots of cell lengths derived from cells grown as in (A). Mean cell length is indicated for each mutant-expressing strain (S.E.M. is indicated) (n ≥ 200 cells for all strains).

Finally, since FtsA(S84L) and FtsA(E14R) are defective for releasing from SUV’s in vitro, including with EDTA, we further tested if Gfp-FtsA(S84L) and Gfp-FtsA(E14R) demonstrate any visible membrane localization in vivo. We analyzed large populations of cells for Gfp-FtsA (wild type or mutant protein) localization and also stained the cells with the membrane dye FM 4-64. We failed to observe colocalization of either Gfp-FtsA(E14R) or Gfp-FtsA(S84L) with the bacterial membrane, and instead observed that both proteins were cytoplasmic, similar to Gfp without FtsA (Fig. S4), and, as reported above, incapable of localizing to a division ring.

## Discussion

Here, we report that FtsA mutant proteins with amino acid substitutions near the site of magnesium coordination in the predicted active site of *E. coli* FtsA (Fig. 1A and 1B) are associated with defects in supporting division in vivo, as well as defects in a variety of in vitro activities. Notably, the degree of the ATP hydrolysis rate defect by each mutant protein does not closely correlate with the severity of functional defects in vivo (i.e., recruitment to division rings or restoring growth at the restrictive temperature in temperature sensitive strains) (Fig. 1C, 1D, and 6A). Consistent with this, FtsA(A188V) does not hydrolyze ATP more rapidly at the permissive temperature relative to the restrictive temperature (Fig. S2), suggesting that temperature sensitive defects mapping to *ftsA* mutations near the nucleotide binding site (Herricks et al., 2014) are suppressed by perhaps another division protein interaction affected by temperature downstream of FtsA. One possibility is that since *ftsA* temperature sensitive mutations are suppressed by changes to the FtsA protomer-protomer interface (Herricks et al., 2014), it is likely a protein that binds to FtsA in a conformation-dependent manner (since polymerization is ATP-dependent) that overcomes *ftsA* defects at low but not high temperature. Consistent with this, temperature-sensitive defects have also been mapped to *ftsK, ftsQ*, and *ftsI*, which can be overcome by FtsN multicopy expression, suggesting that FtsN recruitment and assembly may be the step responsive to thermal-sensitive regulation (Goehring et al., 2007). FtsN also binds directly to FtsA (Baranova et al., 2020).

Here, the in vitro defect that most closely correlates with loss of function in this study appears to be failure to promote ATP-dependent PL remodeling and tubulation, which is a measure of polymerization of FtsA on the PL surface. It is expected that nucleotide binding would promote polymerization by FtsA, and that hydrolysis, supported by an active site magnesium, would regulate how static or dynamic the polymers are by impacting nucleotide occupancy and turnover. Consistent with this, we previously reported that an FtsA variant, without the C-terminal membrane targeting sequence (MTS), forms stable polymers in vitro and that stable FtsA polymers bind to stable FtsZ polymers (Morrison et al., 2022). This suggests that the ability of FtsA to oligomerize into polymers of undetermined length is the critical feature for division that enables association of FtsA with FtsZ at the ring.

Based on the results reported here, purified wild type FtsA appears to be in an equilibrium between the PL-associated and unassociated states (Fig. 2B and 2C), which can be shifted in response to ATP and magnesium, with amino acid residues in the magnesium proximal region being important determinants for the equilibrium, likely by stabilizing one or more conformations. Notably, FtsA(S84L) and FtsA(E14R) fail to polymerize to tubulate vesicles in light scattering assays and by electron microscopy, although they bind to PL’s (Fig. 2B and 2C) and ATP (Fig. 2A), suggesting that Ser 84 and Glu 14 stabilize a conformation relevant for division. Conversely, FtsA(A188V) is defective for binding to SUV, relative to wild type FtsA, which may contribute to the slow rate of ATP hydrolysis observed (Fig. 2A, 2B and 2C). Currently, significant structural information demonstrating various FtsA conformations is lacking; however, conformational flexibility and transitions largely drive differences in actin architecture (i.e., G-actin and F-actin), ATP hydrolysis rates, and protein-protein interactions. It is likely that the diverse array of biochemical and biophysical FtsA activities are all coordinately regulated with polymerization and protein-protein interactions, and that more needs to be understood about FtsA conformations to establish a detailed molecular model of FtsA’s precise role during division.

## Author Contributions

J.J.M., E.M.S., and J.L.C. designed the study, J.J.M., C.N.F., E.M.S., K.N., C.E.T., and B.P. performed experiments, J.J.M., C.N.F., C.E.T., and B.P. edited the manuscript, J.J.M. and J.L.C. wrote the paper, J.L.C. secured funding for the study.

## Acknowledgements

We thank Janet Atoyan for sequencing at the Rhode Island Genomics and Sequencing Center, supported in part by the National Science Foundation (MRI Grant No. DBI-0215393 and EPSCoR Grant No. 0554548 & EPS-1004057), the US Department of Agriculture (Grant Nos. 2002-34438-12688, 2003-34438-13111, and 2008-34438-19246), and the University of Rhode Island. The TEM data was acquired at the RI Consortium for Nanoscience and Nanotechnology, a URI College of Engineering core facility partially funded by the National Science Foundation EPSCoR, Cooperative Agreement #OIA-1655221.

## Supplementary Material

This article contains supporting information.

