## Supplementary Material for "Active site mutations in the bacterial actin homolog FtsA impair direct interactions with the divisome in *Escherichia coli*"

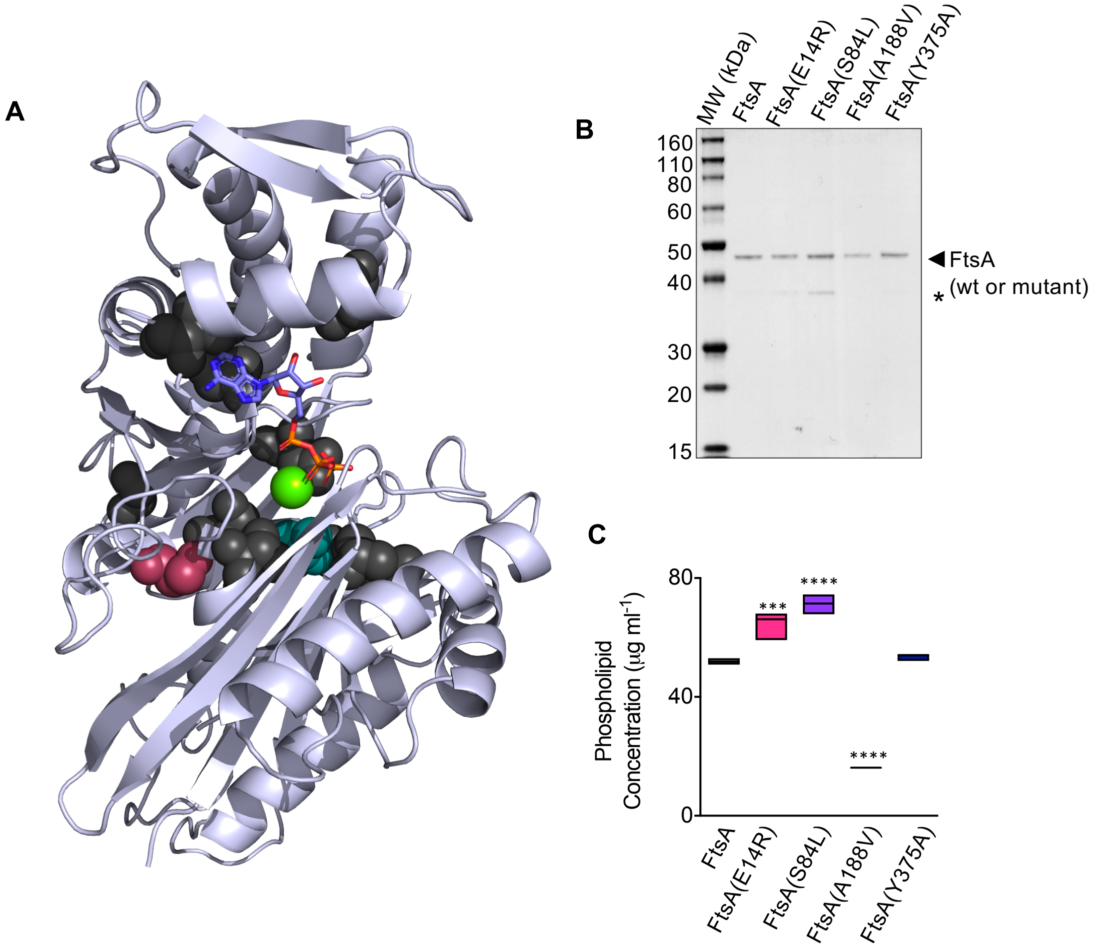


**Figure S1. FtsA wild type and mutant proteins.** (A) *E. coli* FtsA (residues 6-387) modeled onto *T. maritima* FtsA complexed with magnesium [green CPK] and ATP (stick) (pdb: 1E4G). Amino acids associated with mutant *ftsA* alleles that confer temperature sensitive phenotypes are indicated as CPK model (gray), including L83F, C90W, S192L, G205S, D217N, T240I, A338T, I341N, and T378M, in addition to A188V (teal) and S195P (mauve) (Herricks et al., 2014). (B) SDS-PAGE of purified FtsA wild type and mutant proteins used in the study (4 µM). A minor protein contaminant detected in fractions of FtsA wild type and mutant proteins was identified by N-terminal sequencing to be processed OmpF (*). (C) Phospholipid quantitation in fractions of FtsA wild type and mutant proteins (1 µM). Phospholipid concentrations were determined by measuring fluorescence of FM4-64FX membrane dye (1.25 µg $ml^{-1}$) (excitation 555 nm and emission 755 nm) with an Agilent Eclipse spectrofluorometer and comparison to a phospholipid standard curve. Data is representative of at least 3 replicates and error is reported as standard error. Statistical analyses for data in (C) performed by comparison to 'FtsA' (***, p-value = 0.0007; ****, p-value < 0.0001).

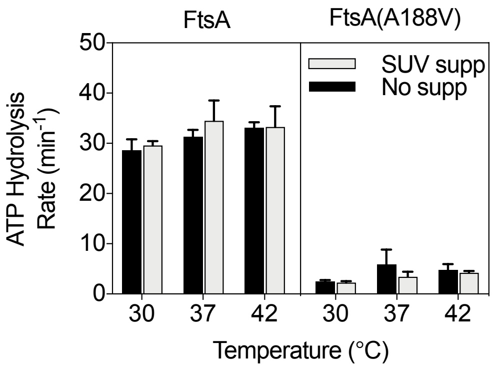


**Fig S2.** Rate of ATP hydrolysis by FtsA (1 µM) or FtsA (A188V) (1 µM) over a range of temperatures and supplemented with SUV's (250 µg ml$^{-1}$) where indicated, as described in *Materials and Methods.*

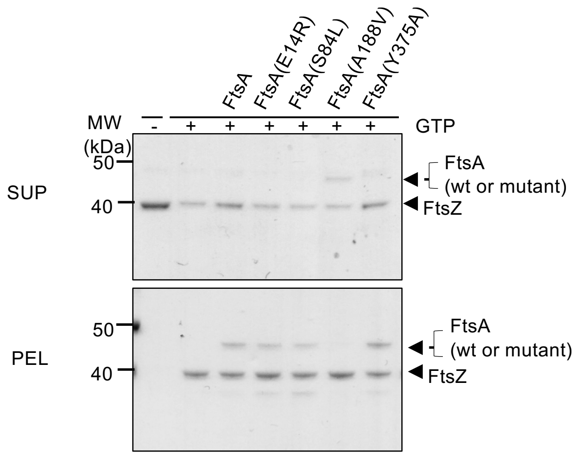


**Fig. S3. Destabilization of FtsZ polymers.** Reactions containing FtsZ (3 μM) and, where indicated, GTP (2 mM), FtsA, FtsA(E14R), FtsA(S84L), FtsA(A188V), FtsA(Y375A) (3 μM), and with ATP (4 mM) and a regenerating system containing acetyl phosphate (15 mM) and acetate kinase (25 μg $ml^{-1}$) were incubated for 5 min then fractionated by ultracentrifugation. Pellets and supernatants were visualized by SDS-PAGE and Coomassie staining and quantified by densitometry. Data is representative of at least 3 replicates.

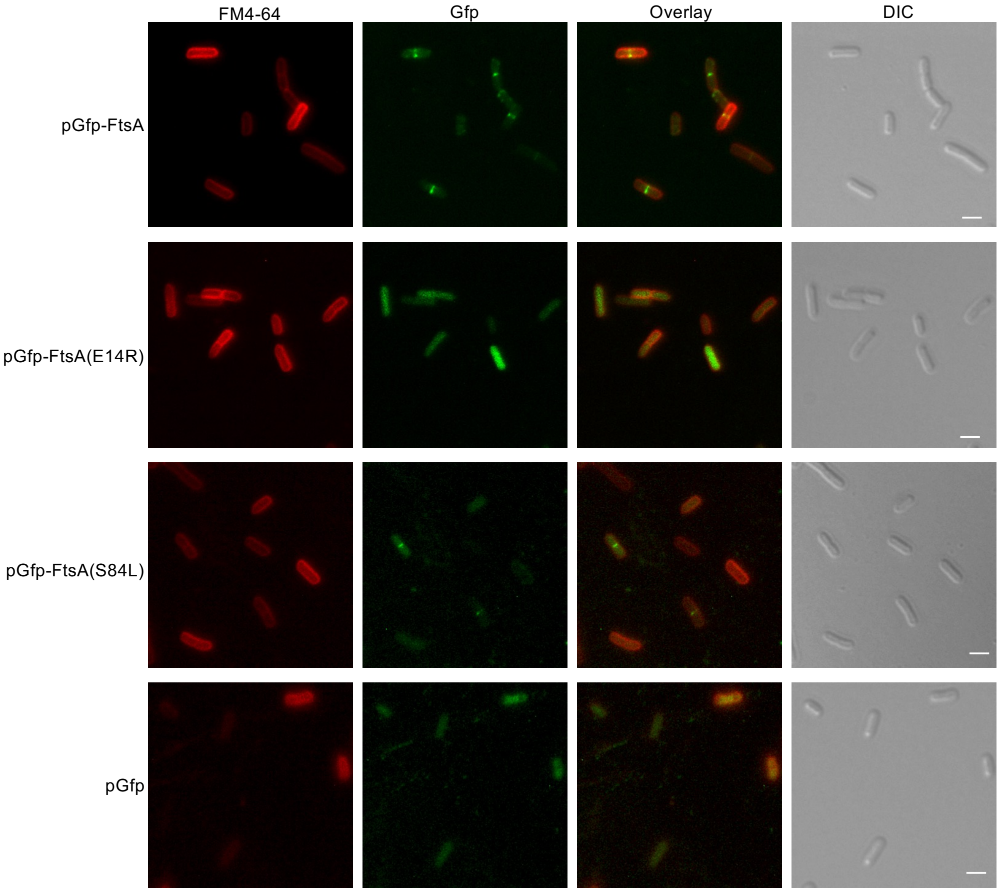


**Fig. S4. Localization and membrane staining of Gfp-FtsA, Gfp-FtsA(E14R) and Gfp-FtsA(S84L).** Fluorescence and DIC microscopy of *E. coli* MG1655 *araE$_{CP}$* cells expressing plasmid-encoded Gfp-FtsA, Gfp-FtsA(E14R), or Gfp-FtsA (Y375A). Cells were grown as described in *Materials and Methods* and stained with FM 4-64FX (3 µg ml$^{-1}$). Scale bars are 2 µm.
